# Time and temperature-resolved transcriptomic analysis of *Arabidopsis* splicing-related mutants

**DOI:** 10.1101/2024.11.08.622454

**Authors:** Sarah Muniz Nardeli, Nabila El Arbi, Varvara Dikaya, Nelson Rojas-Murcia, Daniela Goretti, Markus Schmid

## Abstract

Temperature plays a crucial role in plant growth and development, influencing numerous physiological processes throughout the plant life cycle. Ambient temperature fluctuations can significantly affect transcriptomic adjustments, which are essential for plants to adapt to ever-changing environmental conditions. Despite the known impacts of extreme temperatures on plant physiology, there remains a knowledge gap regarding the specific effects of moderate changes in ambient temperatures on transcriptomic responses. This study employs strand-specific mRNA sequencing (RNA-seq) to assess how different splicing-related mutants respond to varying ambient temperatures, providing a valuable resource to the research community. Analysis of our time-resolved temperature-regulated alternative RNA splicing data reveals that common and exclusive use of the splicing machinery plays pivotal roles in thermoresponsive growth. Furthermore, our analyses demonstrate that moderate temperature changes are translated into widespread transcriptomic responses, including adjustments of the circadian clock and significant splicing changes in light and temperature genes. These results highlight the importance of these particular signaling pathways in adapting to new temperature regimes and suggest future experiments to study the role of alternative RNA splicing in temperature adaptation. Taken together, our results provide insights regarding the role of RNA splicing in plant responses to ambient temperature changes, highlighting the biological relevance of transcriptomic adjustments in enhancing plant resilience and adaptation to climate variability.

**SIGNIFICANCE STATEMENT:**

– This is the first comprehensive study on how mutants involved in multiple steps of the splicing process modulate splicing activity in response to low and high ambient temperature changes.
– We assessed early and acclimated transcriptomic responses and created a valuable resource to investigate the biological outputs.

## INTRODUCTION

Temperature sensing is a vital mechanism that plants use to adjust their development and growth to the seasonal and daily temperature fluctuations that they, as sessile organisms, cannot escape. The molecular processes that enable plants to adapt and adjust their phenotype to varying temperatures are partially known and include diverse mechanisms and signaling pathways, such as adjustments in membrane fluidity, Ca+ signaling, RNA folding, and hormonal changes (Quint et al. 2016; Kerbler and Wigge 2023; Casal et al. 2024). All these changes are largely mediated by transcriptomic adjustments, particularly through RNA splicing, which results in different or no protein products.

In plants, mRNA splicing control is an essential mechanism for plant survival and development. Splicing activity occurs largely co-transcriptionally, which allows plants to regulate the outcome of their mature transcripts and the stoichiometry of their mRNA isoforms quickly when the environment changes. This way, plants can include or exclude sequences that have regulatory roles and, for example, govern transcript stability, localization, structure, interaction with RNA-binding proteins, and more (Sharma et al. 2023). In *Arabidopsis*, splicing is performed by the spliceosome, a multisubunit complex that is well-conserved among eukaryotes, and its function and regulation is essential for embryo and plant development but also plant adaptation under different exogenous stimuli (Kalyna et al. 2012; Marquez et al. 2012; Reddy et al. 2013; Staiger and Brown 2013). The core of the spliceosome is composed of small nuclear ribonucleoproteins (snRNPs), consisting of Sm/LSm proteins (SmB/B′, SmD1, SmD2, SmE, SmF, and SmG; LSm2-8) that form a heptameric ring structure and that associates with the U-rich small nuclear RNAs (snRNAs) (Friesen and Dreyfuss 2000; Golisz et al. 2013). The assembly of snRNPs occurs in the cytoplasm prior to their import into the nucleus, a process that is facilitated by the spliceosome assembly factor GEMIN2 (Schlaen et al. 2015), which is the sole remnant of the mammalian SMN complex, as well as by PRMT5 (Deng et al. 2010; Sanchez et al. 2010) and PICLN (Mateos et al. 2023).

Through the use of alternative splice sites and different exon-intron combinations, alternative splicing (AS) of nascent mRNA transcripts can result in different functional transcripts or non-productive transcript isoforms. While non-functional transcripts with a premature stop codon (PTC) are often targeted for nonsense-mediated decay (NMD), functional transcripts enlarge the gene expression landscape in eukaryotic organisms (Lee and Rio 2015). Intron retention predominates in the majority of intron-containing genes in plants that undergo AS events, as opposed to exon skipping in animals (Marquez et al. 2012). Notably, nuclear-retained, polyadenylated, incompletely spliced transcripts can be stored in the nucleus, ready for splicing activation by environmental or developmental cues (Jia et al. 2020).

AS serves as a critical regulatory mechanism involved in various biological processes, including circadian rhythm regulation and light perception (Sanchez et al. 2010), as well as the transition from vegetative to reproductive phases (Capovilla et al. 2017). Furthermore, AS plays a vital role in enhancing the adaptive capacity of plants when confronted with environmental stresses, such as salinity (Ding et al. 2014) and defense against pathogens (Zhang et al. 2014). Consequently, AS is a fundamental regulatory process in plants, enabling them to modulate their responses in a cell-type-specific, stimuli-specific, or genotype-specific manner (Martín et al. 2021).

In addition, plants shape their transcriptome and use alternative processing of their mRNAs to cope with extreme temperature conditions, such as cold and heat (Calixto et al. 2018, 2019; Ling et al. 2018). In contrast, the effect of moderate ambient temperature changes in mRNA splicing remains largely unexplored, even though recent studies investigated the significance of splicing-associated proteins in modulating gene expression in response to temperature fluctuations and their role in plant adaptation to environmental changes (Schlaen et al. 2015; Capovilla et al. 2018; Huertas et al. 2019; Mateos et al. 2023). However, despite the advancements in how plants respond to low and high ambient temperature variations (Verhage et al., 2017), there remains a knowledge gap regarding the specific responses, particularly in the context of RNA splicing. To address this problem, we used strand-specific RNA sequencing to map the changes in RNA-splicing in response to a shift in temperature over time in *Arabidopsis thaliana* wildtype plants and four mutant genotypes defective in either snRNP assembly or function. Transcriptiome-wide analyses demonstrate the critical role of AS in adjusting key cellular processes, such as circadian clock regulation and light signaling pathways, in response to temperature changes and provide a valuable resource to the scientific community.

## RESULTS AND DISCUSSION

### Splicing-related mutants exhibit general and specific functions in ambient temperature response

To date, it remains largely unexplored how plants respond to moderate changes in ambient temperature. To evaluate whether and how mRNA splicing is impaired at low and high ambient temperatures, we grew *Arabidopsis thaliana* L. (Heyn) wild-type (Col-0) seedlings, core splicing (*pcp-1,smd3b-1*) and assembling (*gemin2-2, prmt5-1*) mutants at continuous low (16°C), optimal (23°C), and high ambient temperatures (27°C) to assess the phenotypic differences in their development and growth. PCP, SMD3b, GEMIN2, and PRMT5 have been shown to be regulated by temperature and to contribute to the adaptive response of plants to temperature variation (Deng et al. 2010; Sanchez et al. 2010) (Schlaen et al. 2015; Capovilla et al. 2018; Huertas et al. 2019; Hong et al. 2023). We found that adult mutant plants exposed to high ambient temperature (27°C) generally showed petiole and leaf elongation similarly to the wild-type. At low ambient temperatures (16°C), all mutant genotypes exhibited a smaller size and pronounced defects, confirming the temperature-dependent developmental differences across genotypes (**Figure 1a**).

**Figure 1.**
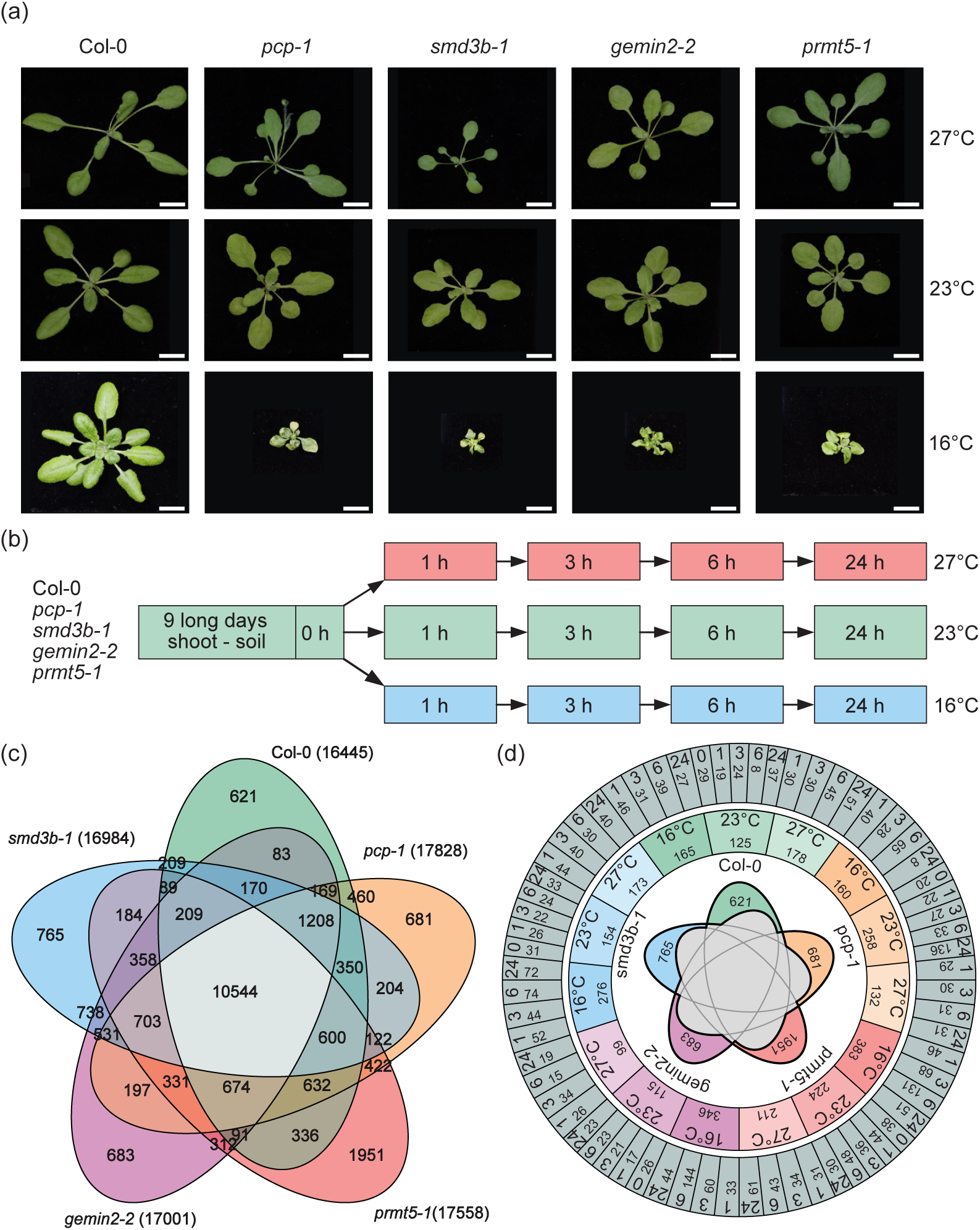
*Arabidopsis* splicing-related mutants in response to different ambient temperatures. (a) Temperature-sensitive phenotypes of *Arabidopsis* wild-type (Col-0) and splicing-related mutants (*pcp-1*, *smd3b-1*, *gemin2-2*, *prmt5-1*) grown continuously at three ambient temperatures (16°C for 31 days; 23°C and 27°C for 19 days). (b) Experimental design for the Ambient Temperature Splicing Transcriptome analysis. (c) Venn diagram showing the overlap of splicing events (PSI) for each genotype across all conditions and time-points. (d) Diagram displaying the number of genotype-specific splicing events divided by ambient temperature (16°C, 23°C, and 27°C) and time of exposure (t0h, t1h, t3h, t6h, t24h). Different colors represent the distinct genotypes in (c) and (d).

To explore the plant responses upon temperature change, we accessed the transcriptome of nine-day-old *Arabidopsis* wild-type seedlings and the four splicing-related mutants across three ambient temperatures (16°C, 23°C, and 27°C) during the course of 24 hours. To avoid false results from comparing different developmental stages, all genotypes were grown under controlled conditions at 23°C for nine days. At this age, wild-type and mutant seedlings did not exhibit any evident macroscopic developmental defects, suggesting that there is no considerable impairment of the spliceosome functions in the young mutant seedlings (**Figure S1**). Samples were collected before the temperature shift (0 hours) and at four distinct time points (1 hour, 3 hours, 6 hours, and 24 hours) post-temperature exposure (**Figure 1b**). The experimental design allowed for a detailed temporal analysis of gene expression and splicing events, with the goal of identifying the early and acclimated transcriptional changes across genotypes. This systematic approach ensured robust comparisons between genotypes and conditions. One hundred ninety-five samples (**Table S1**) were sent to strand-specific mRNA sequencing (**Table S2**). Principal component analysis (PCA) was used to reduce the dimensionality of the gene expression data and visualize the variance between genotypes (**Figure S2**). Overall, the first two principal components explained most of the variance, with a clear separation between the ambient temperatures. For all genotypes, the most distinct transcriptomic responses to temperature occured at 16°C, coinciding with the striking phenotypic defects observed at this temperature in later stages of development, suggesting that gene expression control at this temperature is critical.

Differential gene expression (DE) and differential alternative splicing (DAS) analyses were conducted (**Table S3**), and splicing events were scored for every genotype in all tested conditions (**Table S4**). The overlap of splicing events, calculated as proportion spliced-in (PSI), for all ambient temperatures (16°C, 23°C, and 27°C) and time-points (1-24h) for each genotype is depicted as a Venn diagram (**Figure 1c**). The majority of the splicing events were shared among all genotypes, suggesting that the majority of the splicing events were genotype and temperature-independent. However, a considerable number of events are shared by more than one genotype or presented themselves as a unique response from a particular genotype, with *prmt5-1* displaying the most diverse AS response. We further examined the number of splicing events per genotype at each temperature (**Figure 1d**). This overview of the Ambient Temperature Splicing Transcriptome shows substantial variations in alternative splicing depending on genotype, temperature, and time of the day, underscoring the complexity of temperature-regulated gene expression regulation, with certain mutants exhibiting both general and specific responses.

### Assessment of temperature-dependent AS determines retained intron (RI) as the predominant splicing event type

The number of splicing events, including Alternative 3’ splice site (A3), Alternative 5’ splice site (A5), Skipped Exon (SE), Mutually Exclusive Exons (MX), and Retained Intron (RI), were monitored across different temperatures and time points for the four mutant genotypes (*pcp-1*, *smd3b-1, gemin2-2, prmt5-1*) and compared with the wild-type (Col-0) for each given condition (**Table S5**). The heatmaps illustrate the cumulative number of relative abundances of proportion spliced-in (dPSI) events (**Figure 2**). A normalized scale ranging from 0 to 100% enabled the comparison of the relative distribution of different types of splicing events across all conditions (**Figure 2a**), and identified RI as the most abundant event during the mutant response to ambient temperature changes, in agreement with previous studies showing the prevalence of RI in response to stress in plants (Syed et al. 2012).

**Figure 2.**
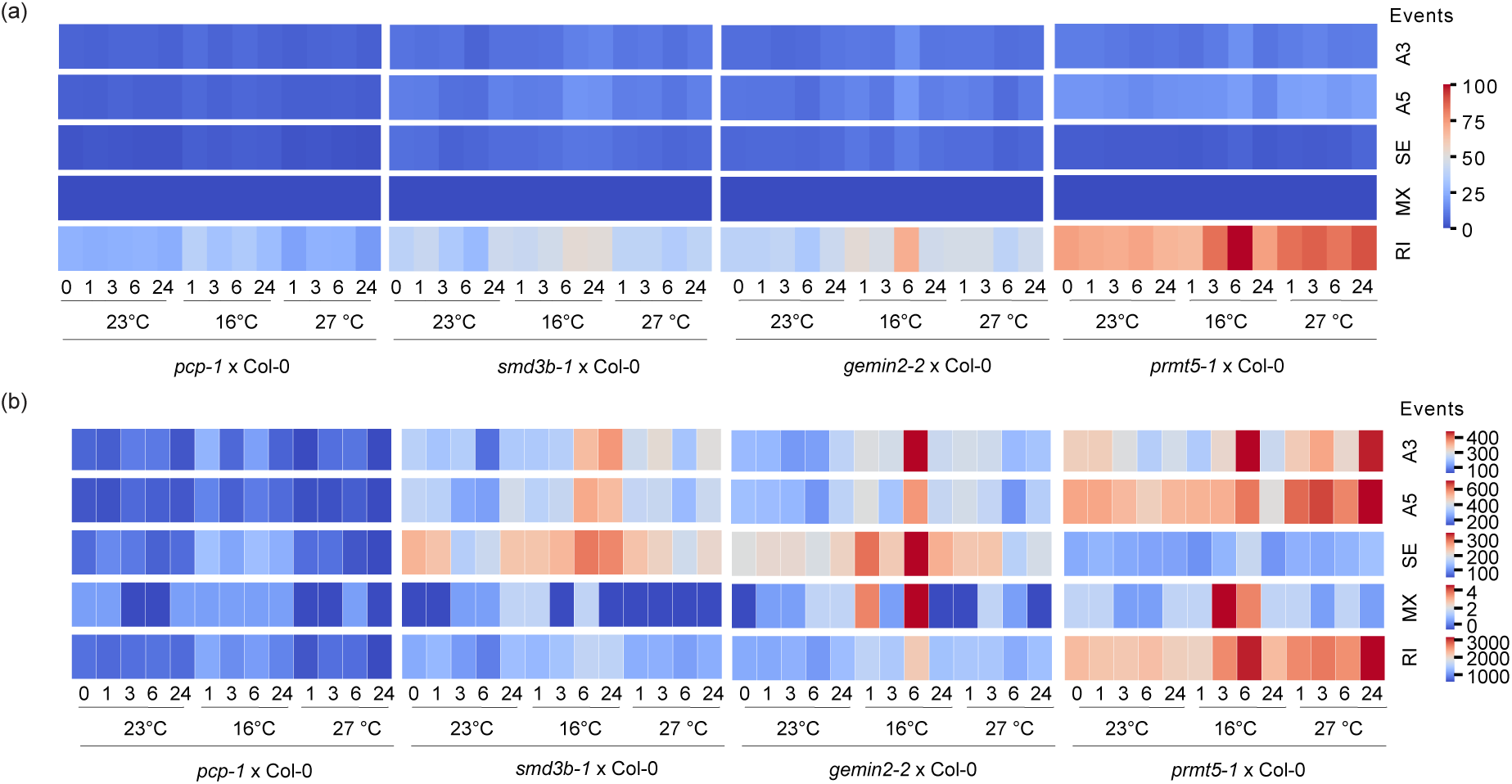
Temperature-dependent alternative splicing events (dPSI) across genotypes and time points. (a) Heatmap representing the relative number of significant alternative splicing (dPSI) events for five splicing types: Alternative 3’ splice site (A3), Alternative 5’ splice site (A5), Skipped Exon (SE), Mutually Exclusive Exons (MX), and Retained Intron (RI) in four genotypes (*pcp-1*, *smd3b-1*, *gemin2-2*, *prmt5-1)* compared to wild-type (Col-0) across three temperatures (16°C, 23°C, and 27°C) at multiple time points (0 hour, 1 hour, 3 hours, 6 hours, and 24 hours) compared to the wild-type Col-0 in the same condition normalized to a maximum of 100 in expression. (b) Heatmap illustrating the absolute number of dPSI events scaled per type of event. The color intensity corresponds to the number of splicing events. Red colors indicate high frequency, as opposed to blue colors for low frequency.

RI transcripts can contain a PTC in their introns, and therefore, they are targeted for degradation through NMD in the cytoplasm; conversely, incompletely spliced, polyadenylated RI transcripts can be retained in the nucleus, and their post-transcriptional splicing can be triggered upon a new developmental or environmental signal (Jia et al. 2020). This expands the gene expression arsenal and serves as a powerful tool to respond to rapid environmental changes, such as ambient temperature fluctuations.

The comparison of the total dPSI number within each type of event (**Figure 2b**) allowed us to better visualize the multitude of different event changes. While it is clear that RI is the prevalent event type, A5, A3, and SE events also play an essential role in the transcriptome adjustment to the new condition, especially in *smd3b-1, gemin2-2*, and *prmt5-1* (Martín et al. 2021). In our study, *pcp-1* clearly presented less differential splicing than the other mutants, which confirms the reported low exon usage in this mutant at 16°C (Capovilla et al. 2018). Interestingly, *prmt5-1* presented a more significant number of different splicing events, especially at high ambient temperatures. It is not surprising that loss of *PRMT5* affects a broader spectrum of AS events since it is involved in multiple aspects of plant development (Deng et al. 2010), and its introns splicing is controlled by various environmental signals (Jia et al. 2020).

### Temperature compensation of the circadian rhythm requires alternative isoforms of the Evening Complex component *LUX*

We noticed that all mutants accumulate more RI events at 16°C, with 6h of exposure being the peak time point for the assembly mutants, *gemin2-2* and *prmt5-1* (**Figure 2**), concordant to the largest number of PSI events (**Figure 1d**). The latter is of particular interest as these two genes have both been previously implicated in the control of the circadian clock (Sanchez et al. 2010; Schlaen et al. 2015), an endogenous oscillator with an approximate period of 24 hours that regulates the timing of cellular processes in most organisms.

In plants, the circadian clock is a complex regulatory system that orchestrates various physiological processes, including flowering time and growth patterns (Dakhiya et al. 2017). In *Arabidopsis,* this clock comprises two primary components: the morning and evening complexes. Each of these complexes consists of specific proteins that play crucial roles in maintaining the circadian rhythm (Hsu and Harmer 2014). The morning complex primarily consists of the core genes *CIRCADIAN CLOCK ASSOCIATED 1* (*CCA1*), *LATE ELONGATED HYPOCOTYL* (*LHY*), and the *PSEUDO-RESPONSE REGULATORS* (*PRR*) including *PRR5, PRR7* and *PRR9*. Conversely, the core evening complex includes *LUX ARRHYTHMO (LUX), TIMING OF CAB EXPRESSION 1* (*TOC1*), *GIGANTEA* (*GI*), and *EARLY FLOWERING GENES 3 and 4* (*ELF3* and *ELF4*) (for details, see **Table 1**).

**Table 1.** Morning and Evening Complex Genes Involved in the Regulation of Circadian Rhythms in *Arabidopsis*.

| Complex | Gene Name | Locus | Description |
| --- | --- | --- | --- |
| Morning Complex | *CCA1 (CIRCADIAN CLOCK ASSOCIATED 1)* | AT2G46830 | A core gene that plays a crucial role in activating morning-phased genes and repressing evening-phased genes. |
|  | *LHY (LATE ELONGATED HYPOCOTYL)* | AT1G01060 | Works alongside CCA1 to regulate the morning genes and repress evening-phased genes. |
|  | *PRR5 (PSEUDO-RESPONSE REGULATOR 5)* | AT5G24470 | Represses morning genes such as CCA1 and LHY, contributing to the daytime circadian rhythm. |
|  | *PRR7 (PSEUDO-RESPONSE REGULATOR 7)* | AT5G02810 | Acts as a repressor of morning-phased genes; interacts with other PRRs to maintain the rhythm. |
|  | *PRR9 (PSEUDO-RESPONSE REGULATOR 9)* | AT2G46790 | Another PRR that suppresses morning-phase genes to regulate daytime responses. |
| Evening Complex | *LUX (LUX ARRHYTHMO)* | AT3G46640 | Part of the evening complex; represses PRR genes during the evening phase and contributes to evening-phase regulation. |
|  | *TOC1 (TIMING OF CAB EXPRESSION 1)* | AT5G61380 | Represses morning genes and regulates evening gene expression; essential for the evening phase of the circadian clock. |
|  | *GI (GIGANTEA)* | AT1G22770 | Regulates flowering time and contributes to the stabilization of evening-phase genes in response to light. |
|  | *ELF3 (EARLY FLOWERING 3)* | AT2G25930 | Works within the evening complex to repress PRR genes and modulate circadian rhythms during the evening. |
|  | *ELF4 (EARLY FLOWERING 4)* | AT2G40080 | Collaborates with ELF3 and LUX to repress PRR genes and regulate evening-phase circadian rhythms. |

Temperature compensation is a critical feature of circadian clocks, allowing organisms to maintain consistent rhythms despite fluctuations in ambient temperature. Alternative splicing is a widespread phenomenon among plant circadian clock genes and a significant portion of AS transcripts in *Arabidopsis* involves RI, which is particularly prevalent in circadian genes (Filichkin and Mockler 2012). Thus, we examined the expression of these core circadian clock-related genes in wild-type seedlings, and *LUX* was the only investigated gene exhibiting a shift in transcript isoform expression in response to our different ambient temperature conditions (**Table S6**).

The temporal expression patterns of the five *LUX* splice variants were assessed in Col-0 at three different temperatures (16°C, 23°C, and 27°C) over a 24-hour period (t0h, t1h, t3h, t6h, t24h) (**Figure 3a**). We observed that *AT3G46640.2* was the major *LUX* isoform expressed at 23°C and also after shifting to 27°C. However, after shifting to low ambient temperature (16°C), *AT3G46640_ID1* became the most abundant isoform, with a gradual induction of the same isoform at high ambient temperature (27°C). Next, we compared the splicing patterns of the three *LUX* predominant isoforms in our conditions, *AT3G46640.2*, *AT3G46640_ID*, and *AT3G46640_P1*, and observed that the temperature-induced dominant ID1 isoform displays an intron retention in the 3’ UTR of the gene (**Figure 3b**).

**Figure 3.**
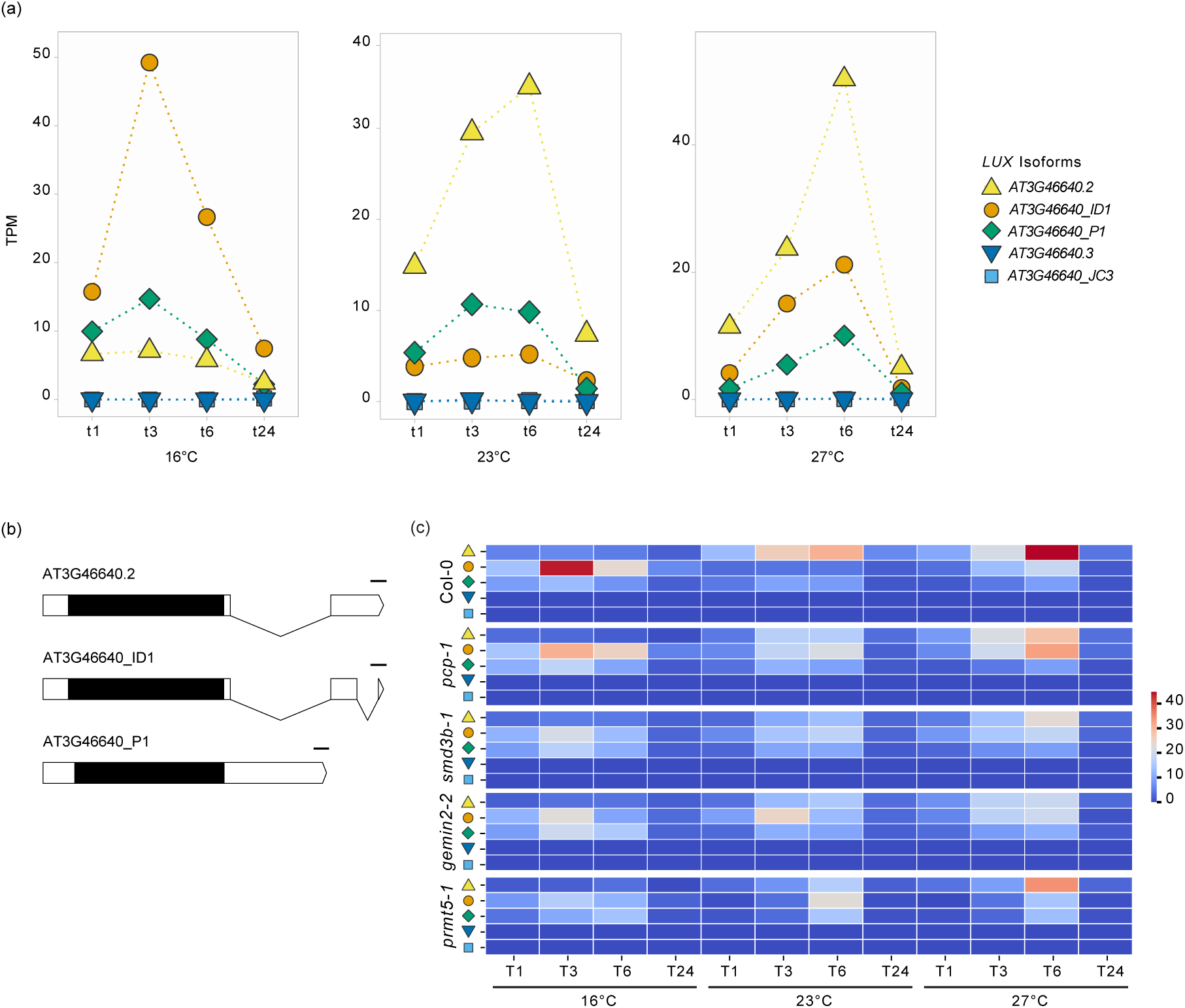
Expression profiles of the circadian clock-associated gene *LUX* under different ambient temperatures. (a) Temporal expression profiles of five *LUX* (AT3G46640) transcripts isoforms for wild-type (Col-0) at three different temperatures (16°C, 23°C, and 27°C) over four time points (t1h, t3h, t6h, and t24h). The expression is presented in TPM (Transcripts Per Million); each colored symbol denotes a different isoform. (b) Transcript structure. Black boxes denote exons, white boxes denote UTRs, and lines denote introns. The bars represent 10 bases. (c) Mean gene expression heatmap of *LUX* splice variants across genotypes, temperatures, and time points. The color intensity in the heatmap indicates the expression (TPM values) of *LUX* isoforms (different colored symbols). The blue-red scale represents TPM values from low (blue) to high (red).

We, therefore, decided to examine the *LUX* expression and AS patterns in the different mutant backgrounds (**Figure 3c, Table S7**). Across all genotypes, expression of the *LUX* splice variants presented modest expression modulation over time. While at optimal ambient temperature (23°C) *AT3G46640.2* was the most abundant isoform in Col-0, there was an increase of the *AT3G46640_ID1* isoform in all mutants. The latter isoform was also the most abundant in the wild-type, *pcp-1,* and *smd3b-1* at 16°C in all time points and in the initial response (up to 3h) in *gemin2-2* and *prmt5-1*. *AT3G46640_P1* isoform increased in expression in the mutants but not in the wild-type in low ambient temperature (**Figure 3c**). We also found that *AT3G46640.2* was the main expressed isoform in Col-0 in high ambient temperature. However, at 27°C, *AT3G46640.2* abundance in the mutants became similar to *AT3G46640_ID1,* and in many time points, they were equally expressed. For detailed gene profiles, see **Figure S3**.

Although LUX AS has not been directly implicated in temperature control, it has been reported that *LUX* is essential for freezing tolerance after cold acclimation, and that the CBF1 transcription factor plays an important role in maintaining *LUX* oscillations in the cold (Chow et al. 2014). Other clock genes have also been reported to undergo alternative splicing when growth temperature changes, with *CCA1* being the best example at low temperatures. The alternative *CCA1* transcript encodes a protein that interferes with the full-length protein, and its suppression is necessary for tolerance at low temperatures (Seo et al. 2012). In contrast to *CCA1*, *LUX* undergoes AS at its 3’ UTR at low temperatures. *LUX* is not the only clock gene with AS at its 3’ UTR. Transcript analysis of the clock genes verified that *LHY*, *LCL1*, and *RVE8/LCL5* mRNAs also have RI events in their 3’ UTRs (Filichkin and Mockler 2012).

The retention of introns in the 3’ UTRs of *LCL1*, *LHY*, and *RVE8* results in transcript variants with abnormally long 3’ UTRs that can be targeted and degraded by NMD, which was further confirmed by *LHY* RI isoform increased in the NMD-impaired *upf1-1* mutant (Filichkin and Mockler 2012). This suggests that similar to these clock genes, RI in *LUX* 3’ UTR may be targeted by NMD to reduce its translational potential and clock-associated activity at suboptimal conditions, such as high and low ambient temperatures. In addition, it is known that elements in plant 3’ UTRs are essential for RNA stability, efficient 3’ end processing (cleavage and poly(A) tail addition), and nuclear export (Xu et al. 2024). This is also consistent with the observation that transcripts with RI isoforms are more likely to be retained in the nucleus under suboptimal light or stress conditions (Jia et al. 2020).

Overall, our data showed that alternative mRNA splicing is regulated by moderate temperature changes, and that an intact spliceosomal machinery is essential for this process since impaired splicing activity could result in disruption of the normal circadian rhythms and efficient AS isoform processing in response to temperature changes.

### Impact of Ambient Temperature on Alternative Splicing of Light and Temperature-Responsive Genes in Splicing-Related Mutants

*gemin2-2* and *prmt5-1* have previously been implicated in the regulation of core clock genes (Sanchez et al., 2010; Schlaen et al., 2015), which is entrained by light input. Our data support the role of thermoresponsive splicing regulation of clock genes across genotypes and we thus extended our analysis to both key light and temperature-related genes.

The clock gene *LUX ARRHYTHMO* (*LUX*) is intricately linked to the regulation of light and temperature-responsive genes, particularly in relation to the *PHYTOCHROME-INTERACTING FACTOR 4* (*PIF4*) (Nusinow et al. 2011), which is a master regulator of temperature sensing and integrates diverse environmental signals (Lucyshyn and Wigge 2009). Hence, to further explore the differential splicing events across genotypes and temperatures, we conducted a comprehensive analysis of several key light and temperature-related genes (**Table 2**).

**Table 2.**
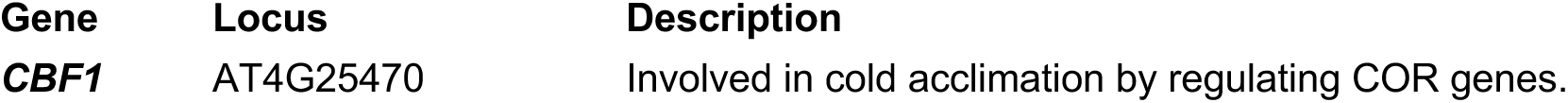
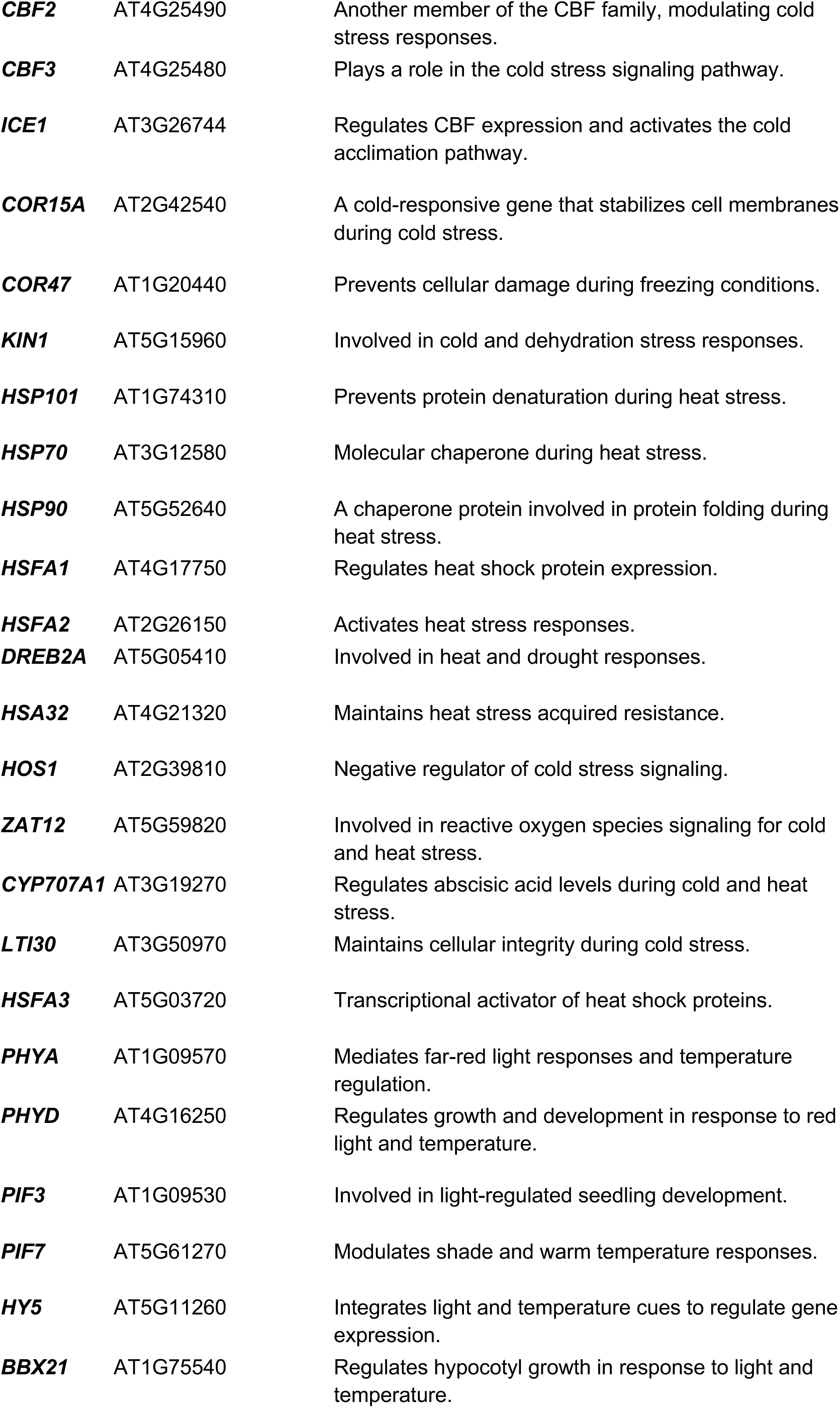
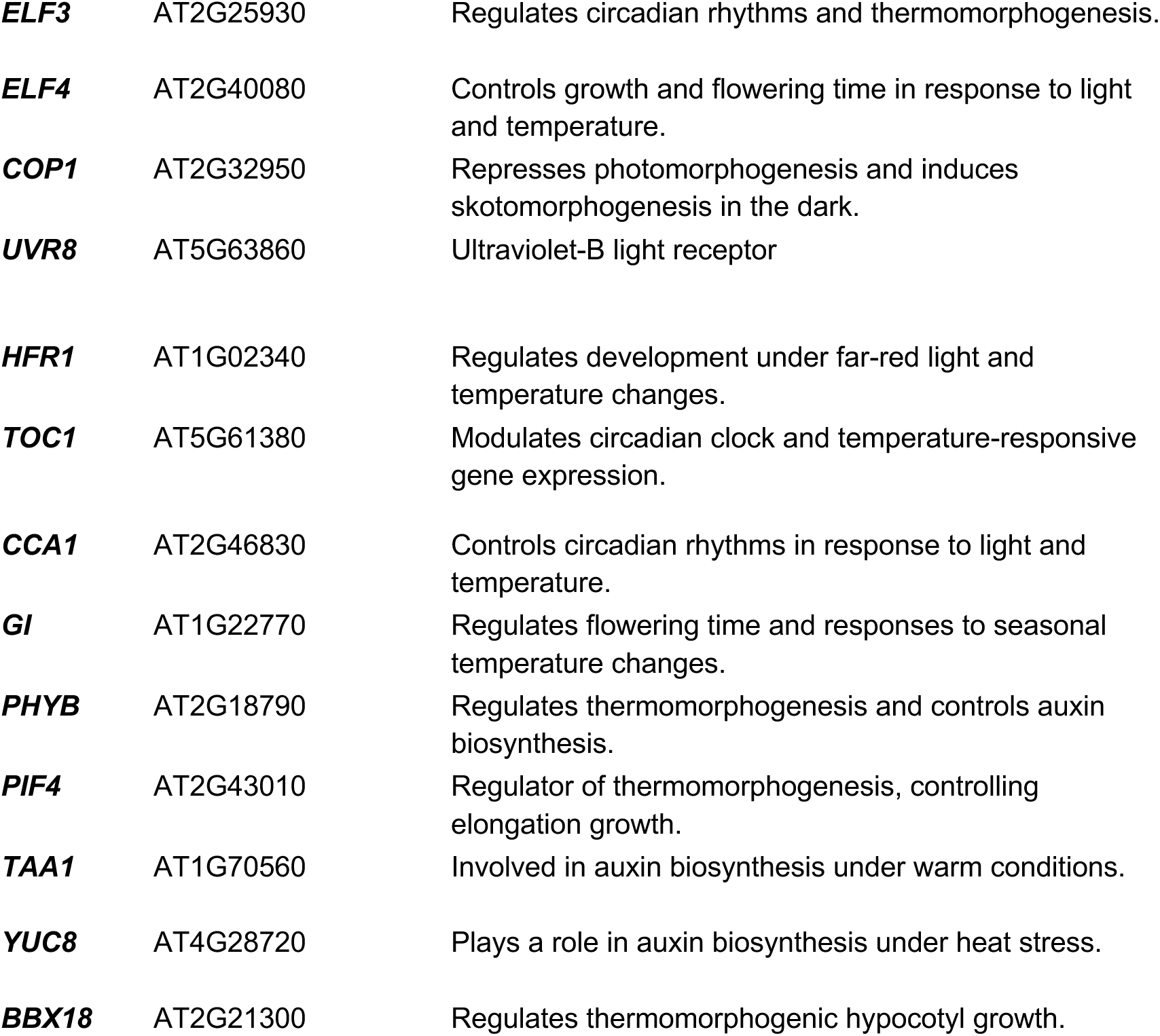
Light and Temperature-Related Genes in *Arabidopsis*.

Of the evaluated genes, nine genes - *PHYB*, *PIF4*, *PIF7*, *HSFA2*, *HSFA3, CCA1*, *TOC1*, *COR15A,* and *ICE1* - presented differential splicing during 24 hours of exposure to different ambient temperatures (**Figure 4**). In all temperatures, a moderate number of dPSI for local events were observed, with *pcp-1* exhibiting fewer splicing changes relative to the other genotypes. At low and high ambient temperatures, all genotypes showed an increase in dPSI (**Figure 4**, **Table S5**). The core clock gene *TOC1* and the light and temperature master regulator *PIF4* presented differential splicing in all conditions. The stress-responsive gene *ICE1* presented more changes in events at all temperatures and genotypes, with the exception of *pcp-1*.

**Figure 4.**
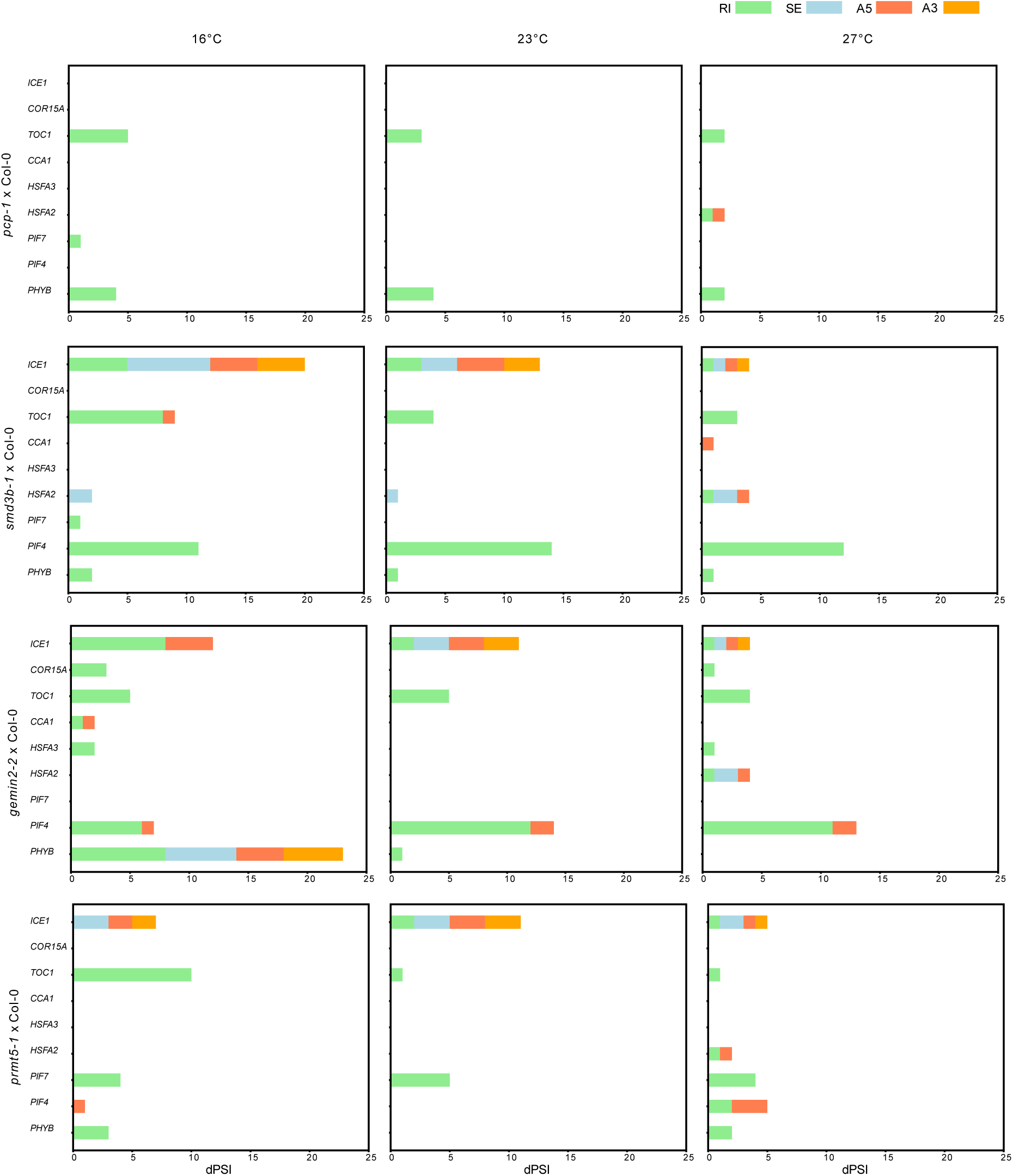
Analysis of difference of Percent Spliced In (dPSI) for Light and Temperature-related genes across different ambient temperatures. Number of differential splicing events with significant dPSI for selected genes across four genotypes (*prmt5-1, smd3b-1, gemin2-2, pcp-1*) compared to the wild-type Col-0 at three temperatures (16°C, 23°C, and 27°C). The different bar colors represent the different types of events: alternative 3’ splice site (A3, golden), alternative 5’ splice site (A5, coral), skipped Exon (SE, light blue), and retained Intron (RI, light green).

*TOC1*, like *LUX*, exhibited temperature-dependent AS, which is vital for its function in maintaining the circadian clock and adapting to environmental changes (James et al. 2012). Here we found that *TOC1* dPSI local events were altered in the splicing-associated mutants, with RI being the most abundant event. However, these changes were not particularly influenced by the temperature shifts in our analysis, rather than the genotype background itself, suggesting that the AS of *TOC1* is mostly influenced by the presence of an integral spliceosomal complex than temperature per se. This does not exclude the possibility that temperature fluctuations may boost this effect, especially at low ambient temperatures (**Figure 4**). In alignment with this, it is reported the potential of *TOC1* to produce distinct AS isoforms under different growth conditions, such as light and temperature, with various regulatory roles (James et al. 2012). *TOC1* and *LUX* are co-regulated and repressed by *CCA1* and *LHY* by binding to the evening element in their promoter region (Alabadí et al. 2001; Tokunaga et al. 2005). In our data, *CCA1* underwent alternative A5 splicing only in the *smd3b-1* mutant at 27°C while in *gemin2-2 CCA1* presented both A5 and RI events only at 16°C. *CCA1* has been reported to display two distinct AS isoforms, *CCA1α* and *CCA1β*, which are oppositely regulated by low or high-temperature change, respectively (Seo et al. 2012). This temperature specificity of *CCA1* AS was also observed in our analysis via AS event count (**Figure 4**). The distinct AS events of *CCA1* seen in *gemin2-2* at 16°C and *smd3b-1* at 27°C further indicate that temperature and specific splicing-related proteins can differentially influence the splicing of this gene. Cold signaling is integrated into the circadian clock through various clock components that may act as upstream regulators of CBFs, such as LHY and CCA1 binding to *CBF* promoters (Dong et al. 2011). *CBF* promoters are also targeted by ICE1 upon cold exposure which induces their expression (Park and Jung 2024). *CBFs* are intronless genes and are known as the central regulators of cold signaling, and upon cold-induction, they bind to specific elements of the promoters of *COR* genes (Ding and Yang 2022), which in turn modulate the downstream physiological changes and cold adaptation. We noticed that *COR15A,* a direct target of *CBFs*, presented RI events at 16°C and 27°C only in *gemin2-2*, while *ICE1* presented a broader variation of AS events (RI, A3, A5, and SE) in all mutants with the exception of *pcp-1*, and throughout temperature range, especially at 16°C (**Figure 4**). Notably, the highest dPSI values for RI events, the main AS event in splicing-associated mutants, was observed at 16°C in *gemin2-2*, which coincides with the *COR15A* RI events, indicating that *GEMIN2* may be important for the ICE1-CBFs-CORs signaling pathway under low temperatures. In addition, the accumulation and variation of AS events recorded for *ICE1* in all the mutants and temperatures support the view that AS is a vital mechanism for fine-tuning responses to environmental stress (Dikaya et al. 2021).

CBF regulation is not restricted only to cold or low-temperature changes. Their expression can also be modulated by PIF7 and PIF4 in a circadian manner (Lee and Thomashow 2012), and PIF7, in turn, is regulated by the clock component TOC1 and PHYB, a well-studied photoreceptor (Kidokoro et al. 2009). PIF4 and PIF7 are master regulators of the high temperature and light quality signaling in plants (Delker et al. 2022). Interestingly, we noticed that *PIF4* presented AS events in all the mutants but *pcp-1*, with RI being the primary and most abundant type across all temperatures (**Figure 4**). *PIF4* also presented differential A5 local events in *smd3b-1* and *prmt5-1* but to a lesser extent. On the other hand, *PIF7* presented alternative RI in all temperatures in *prmt5-1* and, to a lesser extent, in *pcp-1* and *smd3b-1* only at 16°C. However, *phyB* displayed substantial RI events in all the mutants and almost all conditions, except *gemin2-2* at 27°C and *prmt5-1* at 23°C. Strikingly, we observed that *phyB* is under extensive AS with differential local AS type of events present only in *gemin2-2* at 16°C (**Figure 4**). These data suggest that distinct splicing factors contribute uniquely to various components of well-studied signaling pathways, such as the phyB-PIF network.

An increase in growth temperature leads to the re-adjustment of molecular activities required for thermotolerance. Several transcription factors belonging to different protein families are involved in the upregulation of these genes (Baniwal et al. 2004; Fragkostefanakis et al. 2015). Among them, heat stress transcription factors (HSF) are considered indispensable as they control the expression of the protein chaperones (Scharf et al. 2012). Many HSFs undergo AS, especially at the 3’ UTR, generating aberrant transcripts targeted by NMD (Sugio et al. 2009; Liu et al. 2013). Interestingly, *Arabidopsis HSFA2* AS, under severe stress conditions, leads to an mRNA with long 3’ UTR that can still be translated and escape NMD targeting (Liu et al. 2013). Other AS events like RI were also present and can lead to alternative mRNAs with different protein domains. The loss of the splicing-related protein LSM5 results in thermosensitivity, a phenotype partly attributed to the misplicing of *HSFA3* and a DnaJ-domain chaperone protein (Cui et al. 2014; Okamoto et al. 2016). Loss of STABILIZED1 (STA1) splicing factor also causes problems in thermotolerance acquisition via the defective splicing of *HSFA3* and several other HSPs under heat stress (Kim et al. 2017, 2018). We found that *HSFA2* is differentially spliced at 27°C in *pcp-1, gemin2-2*, and *prmt5-1,* and in a temperature-independent manner in *smdb3-1. HSFA3* also underwent AS in low and high ambient temperatures in *gemin2-2* (**Figure 4**). The splicing variations in these heat shock factors suggest a complex regulatory network that enables plants to optimize stress responses through post-transcriptional modifications.

It was previously proposed that temperature directly influences alternative splicing of genes, and that this altered splicing machinery affects the splicing of downstream genes that have a role in temperature sensing (Verhage et al. 2017). Our study of splicing mutants responding to temperature fluctuations is a key tool in understanding temperature-regulated AS. Despite the macroscopic phenotypic similarities between wild-type and splicing-related mutants at the early stages of development, the transcriptomic adjustments are mutant-specific, as they display distinct gene expression, with great contribution of alternative splicing patterns. the plant’s growth and development.

## EXPERIMENTAL PROCEDURES

### Plant material

Wild-type (Col-0) and the splicing mutants *pcp-1* (SALK_089521), *smd3b-1* (SALK_006410), *gemin2-2* (SAIL_567_D05), and *prmt5-1* (SALK_065814) were grown in soil under long-day conditions in Percival chambers (16 h light/8 h dark, 120 μmol m⁻² s⁻¹ full-spectrum white light LEDs, 65% relative humidity) for nine days at 23°C (0 h, zeitgeber time six) followed by a shift to 16°C, 27°C, or maintenance at 23°C for 1 h, 3 h, 6 h, and 24 h. For each time point, the aerial parts of the seedlings were harvested in liquid nitrogen, and the plant material was stored in ultra-freezers. For each condition, a minimum of 15 seedlings and three biological replicates were harvested.

### Sample preparation and RNA extraction

Total RNA from 195 samples (wildtype and four splicing mutants, see Table S1) was isolated using the RNeasy Plant Mini Kit (Qiagen, Hilden, Germany), followed by Turbo DNAse treatment (Thermo Fisher Scientific, Waltham, MA, USA) according to the manufacturer’s guidelines. RNA concentration was measured using a Nanodrop 1000 (Thermo Fisher Scientific, Waltham, MA, USA) and stored at −70°C until further use.

### Data generation

High-quality RNA samples were sent to Novogene (UK), where the RNA Integrity Number (RIN) was assessed using a 2100 Bioanalyzer with the RNA 6000 Nano kit (Agilent, Santa Clara, USA). Strand-specific mRNA libraries were prepared with poly-A enrichment and sequenced on the NovaSeq 6000 platform (Illumina, San Diego, USA) with 150-bp paired-end reads to an average depth of 50 million read pairs (see **Table S2**).

### RNA-sequencing data processing

Data was pre-processed using nf-core/rnaseq v3.12.0 (Patel et al. 2024) of the nf-core collection of workflows (Ewels *et al*., 2020) utilizing reproducible software environments from the Bioconda (Grüning *et al*., 2018) and Biocontainers (da Veiga Leprevost et al., 2017) projects. The pipeline was executed with Nextflow v23.10.1 (Di Tommaso *et al*., 2017) with the following command:

[nextflow run nf-core/rnaseq -r 3.12.0 -profile uppmax --outdir rnaseq_AtRTD2-QUASI --project naiss2023-22-795 --input samplesheet_rnaseq_ csv --skip_alignment --pseudo_aligner salmon --fasta AtRTD2_QUASI_gentrome.fa --gtf AtRTDv2_QUASI_19April2016.gtf --salmon_index AtRTD2_QUASI_gentrome_salmon_v1dot4dot0].

Briefly, the quality of raw reads was assessed using FastQC (v0.11.9); over-represented sequences and residual adapter sequences were trimmed from raw reads using Cutadapt (v3.4) (Martin 2011) and Trimgalore (v0.6.7) (Krueger et al. 2023) with default settings. After trimming, only paired-end reads were used for further downstream analysis. Paired-end reads were aligned to the *Arabidopsis thaliana* transcriptome (AtRTD2-QUASI) (Zhang et al. 2017) using Salmon (v2.6.0) (Patro et al. 2017). Post-processing and analysis of the RNA sequencing data were performed using the 3D RNA sequencing app (Guo et al. 2020). TMM (weighted trimmed mean of M-values) was used as the data normalization method, and the output data were filtered for an absolute log2 fold change (log2FC) of 0.5 or higher (see **Table S3**).

### Alternative splicing analysis

To calculate local alternative splicing events based on the expression of transcripts in each dataset, we utilized SUPPA2 (Trincado et al. 2018). Data were processed using nf-core/rnasplice v1.0.1 (Ashmore et al. 2024) from the nf-core collection of workflows (Ewels et al. 2020) utilizing reproducible software environments from the Bioconda (Grüning et al. 2018) and Biocontainers (da Veiga Leprevost et al. 2017) projects. The pipeline was executed with Nextflow v23.10 (Di Tommaso et al. 2017) using the following command:

[nextflow run nf-core/rnasplice -profile uppmax --input samplesheet_195_salmon_results_AtRTD2-QUASI.csv --contrasts contrast_pertimepoint_withingenotype.csv --source salmon_results --skip_alignment --suppa --outdir RNAsplice --fasta ATRTD2-QUASI/AtRTD2_QUASI_gentrome.fa --gtf ATRTD2-QUASI/AtRTDv2_QUASI_19April2016.gtf --salmon_index ATRTD2-QUASI/AtRTD2_QUASI_gentrome_salmon_v1dot4dot0 --project naiss2023-22-795 -bg -r 1.0.1]

SUPPA2 is instrumental in studying splicing at the local alternative splicing level or at the transcript isoform level. We analyzed seven types of alternative splicing events: (1) Alternative 3′ splice site (A3-event), where the 3′ site acts as an acceptor; (2) Alternative 5′ splice site (A5-event), where the 5′ site acts as a donor; (3) Alternative first exon (AF-event), where the first exon is retained after splicing; (4) Alternative last exon (AL-event), where the last exon is retained after splicing; (5) Mutually exclusive event (MX-event), where one of two exons is retained; (6) Retention intron event (RI-event), where the intron is confined within mRNA; and (7) Skipping exon event (SE-event), where an exon may be spliced out or retained. For each gene in a given genotype or condition, the average percent spliced-in (PSI) value was calculated. Alternative splicing of a gene in a particular dataset is considered only if it fulfills the criteria of PSI < 0.9 and > 0.1 (see **Table S4**). Using the PSI cut-off, we detected specific local events that occurred in either the wild-type or the mutant datasets. Additionally, we employed SUPPA2 to conduct differential splicing analysis (dPSI), performing pairwise comparisons between each mutant and wild-type. Differentially spliced candidates were identified using a p-value threshold of < 0.05 (see **Table S5**).

### Heatmaps and Gene Profiles

Custom Python scripts were developed in Jupyter notebooks for generating heatmaps, gene expression profiles, proportion spliced-in (PSI) and differential splicing (dPSI) gain and loss profiles, and intron-exon gene structures using the Matplotlib, Pandas, Numpy, Scipy, and Seaborn packages.

## DATA AVAILABILITY STATEMENT

The datasets analyzed in the current study are available in ENA PRJEB81550. The Experimental Procedures section refers to all freely available programs and packages used in this manuscript.

## ACKNOWLEDGEMENTS

Vasiliki Zacharaki and Thomas Dobrenel for their assistance during the greenhouse experiments. UPSC Bioinformatics facility for data deposit to ENA. The computations and data handling were enabled by resources provided by the National Academic Infrastructure for Supercomputing in Sweden (NAISS), partially funded by the Swedish Research Council through grant agreement no. 2022-06725. This work has been supported by a grant from the Knut och Alice Wallenberg Stiftelse (KAW 2018.0202) to M.S.

## AUTHOR CONTRIBUTIONS

SMN, NEA, VD, NRM, DG: Sample acquisition and preparation. SMN: Data processing, analysis, and visualization. SMN, NEA, MS: Manuscript writing. SMN, NEA, VD, MS figure design. MS: Funding acquisition.

