## Supporting Figures S1 - S3 for "Time and temperature-resolved transcriptomic analysis of *Arabidopsis* splicing-related mutants"

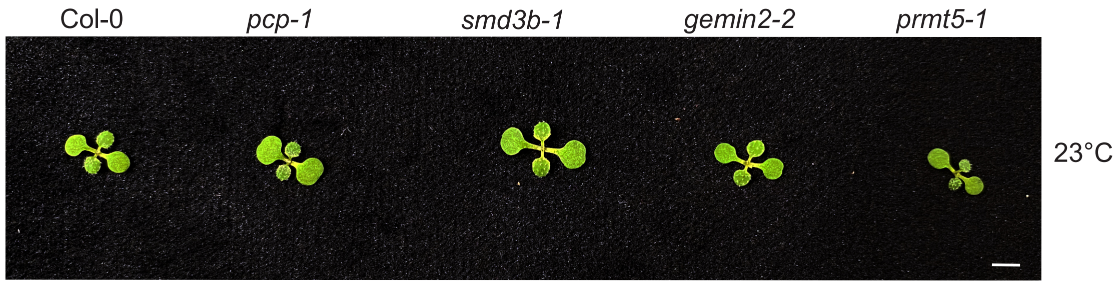


**Figure S1. Splicing-related mutant seedlings in response to different ambient temperatures.** Representative of temperature-sensitive phenotypes of *Arabidopsis* wild-type (Col-0) and splicing-related mutants for genes involved in Sm-ring subunits (*pcp-1* and *smd3b-1*) and snRNP assembly (*gemin2-2* and *prmt5-1*) grown at continuous 23°C per nine days. The scale bar is 0.5 cm.

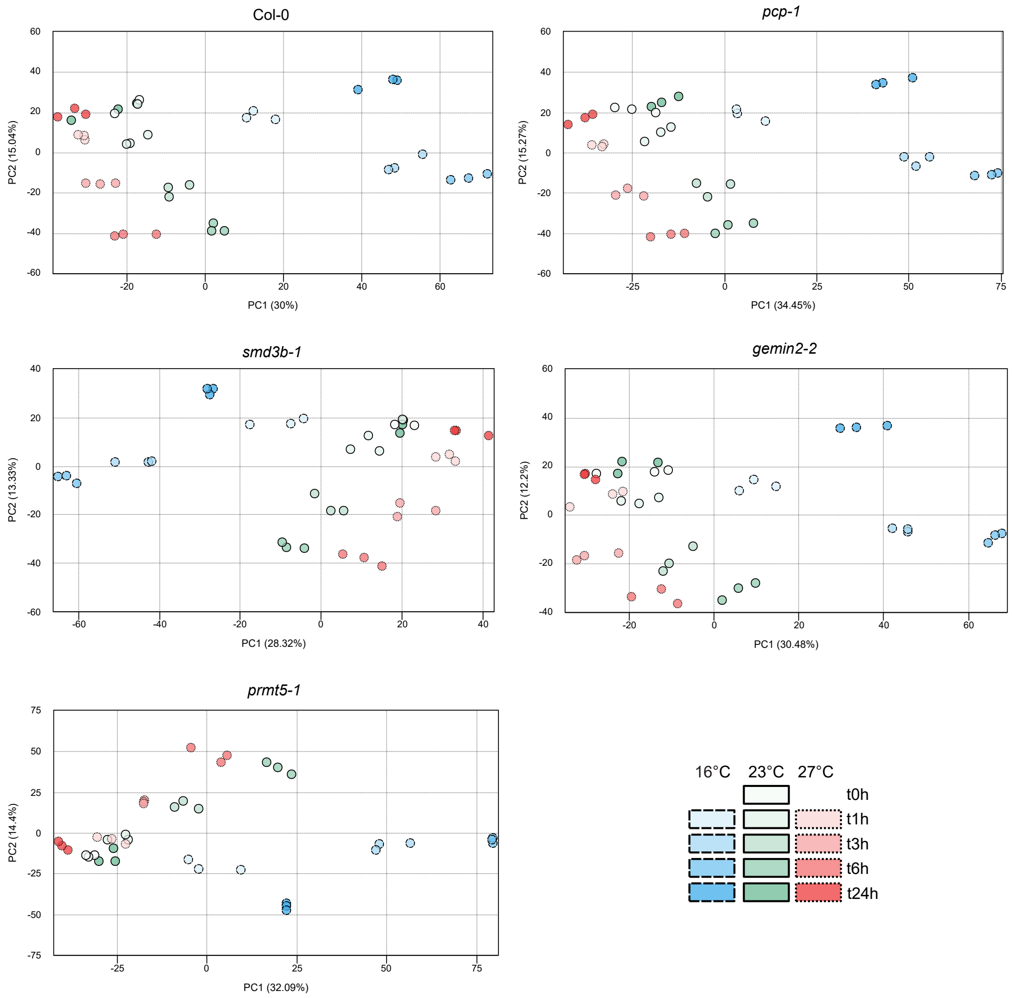


**Figure S2. Principal Component Analysis (PCA) of transcriptomic profiles across genotypes, temperatures, and time-points.** *Arabidopsis* seedlings cultivated for nine days at a constant temperature of 23°C (designated as t0h and solid lines) shifted to 16°C (represented by blue circles and dashed lines) and 27°C (indicated by red circles and dotted lines), with a control group maintained at the original temperature of 23°C (illustrated by green circles). The color intensity of the circles corresponds to the temporal progression of the temperature treatments, with lighter shades indicating shorter exposure times (t1h) and progressively darker shades representing longer exposure durations, culminating in the darkest shade for the 24-hour time point (t24h). The baseline condition at time point 0 hours (t0h) is denoted by a white circle, serving as a reference for the subsequent temperature treatments.

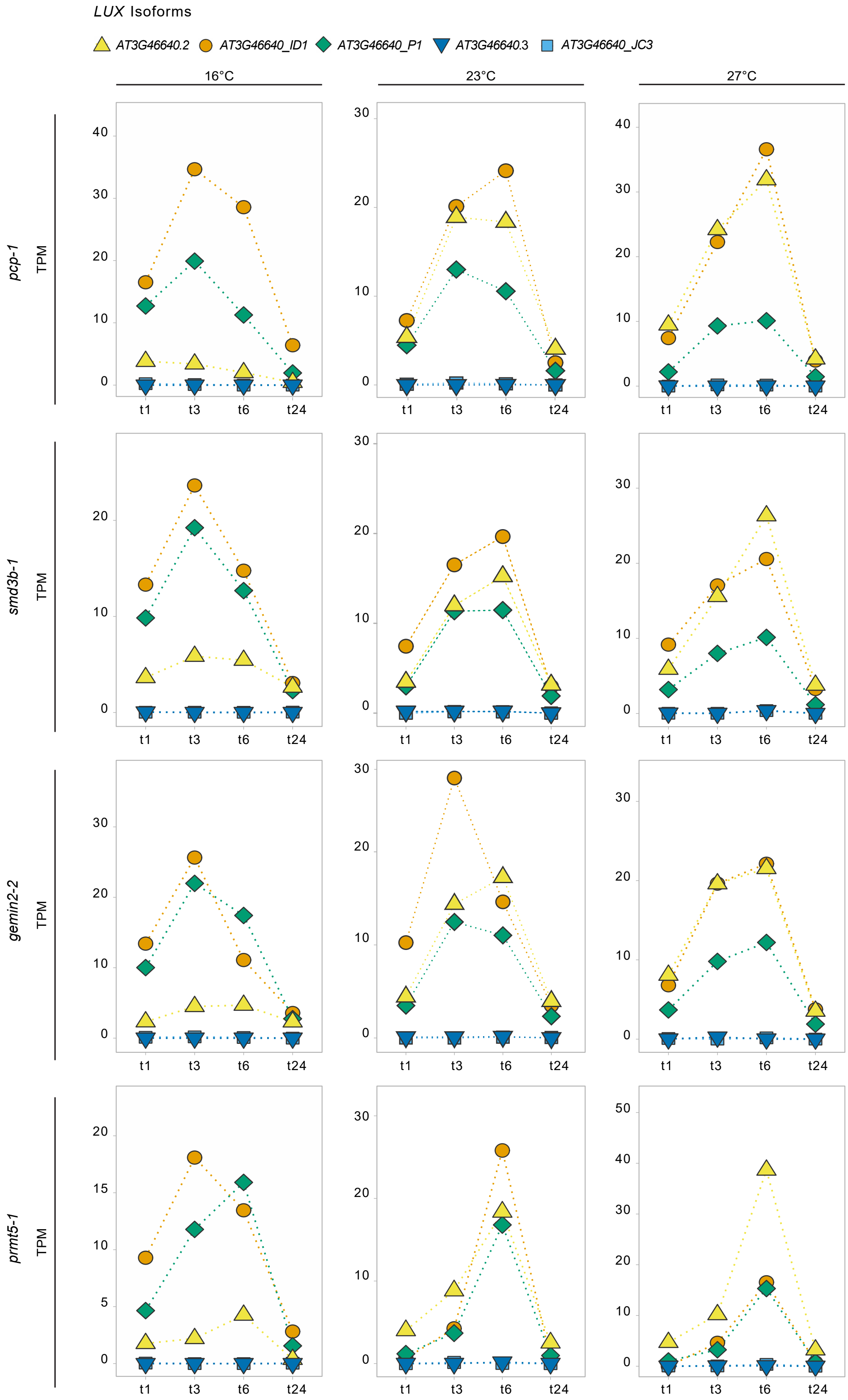


**Figure S3. Temporal expression profiles of *LUX* transcripts in the splicing-related mutants across different temperatures**. The different colored symbols represent the different transcript isoforms. Time-course in the x-axis; TPM values for the different genotypes in the y-axis.
